# Complete chloroplast genomes and phylogeny in three *Euterpe* palms (*E. edulis, E. oleraceae* and *E. precatoria*) from different Brazilian biomes

**DOI:** 10.1101/2022.03.21.485093

**Authors:** Ana Flávia Francisconi, Luiz Augusto Cauz dos Santos, Jonathan Andre Morales Marroquín, Cássio van den Berg, Alessandro Alves-Pereira, Luciano Delmondes de Alencar, Doriane Picanço-Rodrigues, Cesar Augusto Zanello, Marcones Ferreira Costa, Maria Teresa Gomes Lopes, Elizabeth Ann Veasey, Maria Imaculada Zucchi

**Affiliations:** Programa de pós-gradução em Genética e Biologia Molecular Universidade Estadual de Campinas, Campinas, São Paulo, Brasil; Department of Botany and Biodiversity Research, University of Vienna, Wien, Austria; Departamento de Ciências Biológicas, Universidade Estadual de Feira de Santana, Feira de Santana, Bahia, Brasil; Departamento de Genética, Universidade de São Paulo, Piracicaba, São Paulo, Brasil; Departament de Biologia Vegetal, Universidade Estadual de Campinas, Campinas, São Paulo, Brasil; Departamento de Biologia, Universidade Federal do Amazonas (ICB-UFAM), Manaus, Amazonas, Brasil; Universidade Federal do Piauí, Floriano, Piauí, Brasil; Departamento de Produção Animal e Vegetal, Universidade Federal do Amazonas, Manaus, Amazonas, Brasil; Agência Paulista de Tecnologia dos Agronegócios, Piracicaba, São Paulo, Brasil

**Author notes:** Corresponding author (MIZ), (AFF).

**Keywords:** Arecaceae, SNPs, genome-skimming, genome comparison, polymorphism

## Abstract

The Brazilian palm fruits and hearts-of-palm of *Euterpe edulis, E. oleracea and E. precatoria* are an important source for agro-industrial production, but overexploitation requires conservation strategies to maintain genetic diversity. Chloroplast genomes have conserved sequences, which are useful to explore evolutionary questions. Besides the plastid DNA, genome skimming allows the identification of other genomic resources, such as single nucleotide polymorphisms (SNPs), providing information about the genetic diversity of species. We sequenced the chloroplast genome and identified gene content in the three *Euterpe* species. We performed comparative analyses, described the polymorphisms among the chloroplast genome sequences (repeats, indels and SNPs) and performed a phylogenomic inference based on 55 palm species chloroplast genomes. Finally, using the remaining data from genome skimming, the nuclear and mitochondrial reads, we identified SNPs and estimated the genetic diversity among these *Euterpe* species. The *Euterpe* chloroplast genomes varied from 159,232 to 159,275 bp and presented a conserved quadripartite structure with high synteny with other palms. In a pairwise comparison, we found a greater number of insertions/deletions (indels = 93 and 103) and SNPs (284 and 254) between *E. edulis*/*E. oleracea* and *E. edulis*/*E. precatoria* when compared to *E. oleracea*/*E. precatoria* (58 indels and 114 SNPs). Also, the phylogeny indicated a closer relationship between *E. oleracea*/*E. precatoria*. The nuclear and mitochondrial genome analyses identified 1,077 SNPs and high divergence among species (F_ST_ = 0.77), especially between *E. edulis* and *E. precatoria* (F_ST_ = 0.86). These results showed that, despite the few structural differences among the chloroplast genomes of these *Euterpe* palms, a differentiation between *E. edulis* and the other *Euterpe* species can be identified by point mutations. This study not only brings new knowledge about the evolution of *Euterpe* chloroplast genomes, but also these new resources open the way for future phylogenomic inferences and comparative analyses within Arecaceae.

## Introduction

The palm family (Arecaceae) comprises 188 genera and ca. 2,400 species, distributed throughout tropical and subtropical regions [1,2]. Palms are often keystone species, providing ecosystem services, shaping their environment and offering many products used for fabrics, fuel, food, medicine and as ornamentals [2–4]. Within Arecaceae, the genus *Euterpe* originated in South America and includes seven species distributed throughout Central America and tropical South America [2,5,6]. In Brazil, among the native species of *Euterpe, E. edulis* Mart., *E. oleracea* Mart. and *E. precatoria* Mart. are the most important from an agro-industrial point of view [7].

*Euterpe edulis* is endemic to the Atlantic Forest and has substantial importance for the functioning of the ecosystem [8,9]. However, the overexploitation of heart-of-palm [10,11] has placed this palm species in the Brazilian Flora Red Book [12] as “vulnerable (VU)”, which makes conservation strategies for it extremely necessary. Recently, as an alternative to palm heart-of-palm extraction, local communities have started to cultivate this species for fruit pulp extraction, due to the similarity with the Amazonian açaí (*E. oleracea* and *E. precatoria*) [9,10,13]. This practice has been fundamental to capture and conserve the species genetic diversity present in older forests [10].

*Euterpe oleracea* and *E. precatoria* are the main sources for the production of açaí fruit pulp, estimated to be 760,000 tons in 2017, mostly in the northern region of Brazil (98,36% of the national production) [14,15]. *E. oleracea* is a multiple-stemmed palm, commonly found in the Amazon estuary, especially prevalent in the Brazilian states of Pará, Maranhão and Amapá [15–17]. However, in the last 20 years, the intense demand for açaí fruit in Pará resulted in a significant loss of local tree species richness, which presumably is occurring after decades of thinning to reduce interspecific competition with açai palm trees [18]. This resulted in a reduction of pollinators, which in turn decreased the fruit production [19]. *E. precatoria* is a single stemmed palm, predominantly found in the Brazilian states of Amazonas, Acre and Rondônia [20], in non-flooded areas [21]. This species is highly promising for açaí fruit extraction, as the state of Amazonas is already the second largest producer of Brazil [15]. Therefore, the establishment of strategies for *E. precatoria* domestication, conservation and management of populations is extremely necessary, in addition to the support of riverside communities and farmers to conserve this species based on their genetic diversity [15].

The fast growth of açaí (*E. oleracea* and *E. precatoria*) pulp production in the last decades and the threats of extinction for *E. edulis* indicate an urgent demand for new genomic resources in this genus, contributing to the understanding of the evolution and population dynamics of the species, which can result in the best planning of sustainable development and conservation [9,15]. Fortunately, high-throughput sequencing technologies have revolutionized how genomic data can be obtained for plant species and have allowed the complete assembly of the chloroplast genome for an increasing number of non-model species [22–24], while providing useful information for the study of genetic variation of native species.

The common structure of the chloroplast genomes typically has four parts: two single copy regions, one large (LSC) and one small (SSC), and a pair of inverted regions (IRs) [25] which basically consist of long circular or linear molecules (120 to 180 kb), usually conserved across palms species [26]. Apart from the conservative nature and low nucleotide substitution rates of the chloroplast sequences [27], the description of this organellar genome is important to identify repetitive sequences and polymorphic regions, useful to further obtain molecular markers, which can be applied to assess genetic structure and diversity of natural populations [28,29]. Whole chloroplast genomes and chloroplast sequences have been used to infer phylogenetic relationships and investigate diversification patterns [24,30–32]. The tribe Euterpeae was already analyzed with combined chloroplast and nuclear sequence data [33], and whole chloroplast genomes have been reported in reference for two species in the *Euterpe* genus (*E. oleracea* and *E. edulis*) [34]. These priors research emphasize the differentiation of *E. edulis*, due to its distribution in a different Brazilian biome (Atlantic Forest biome). But, the addition of new chloroplast genomes can give more consistency to the evolutionary processes of the genus.

Moreover, the amount of plastid sequences present in a data from total genomic DNA extraction depends on different factors and can vary substantially, from 0.4% to 29.5% [23], and the recovery of these sequences can occur through the use of genome skimming. Therefore, the remaining nuclear and mitochondrial sequences can provide genomic resources [27,35], representing useful information to study the genetic variation of native species.

Considering the effectiveness of genome skimming to produce genomic data, we employed this approach to obtain three newly sequenced complete chloroplast genomes, thereby facilitating the study of genetic diversity from *Euterpe* species. We were able to investigate: i) the plastid organization in *Euterpe* and the synteny level with other available Arecaceae species; ii) the repetitive sequences and polymorphic regions in the chloroplast genomes; iii) the Arecaceae family evolution based on a phylogenomic study with complete chloroplast genomes; and iv) the species divergence using the remaining nuclear and mitochondrial data from genome-skimming.

## Material and Methods

### Leaf material

The leaf material from *Euterpe edulis* and *E. oleracea* were collected from the *ex situ* collection at the Escola Superior de Agricultura “Luiz de Queiroz” (ESALQ), Universidade de São Paulo (USP) in Piracicaba, São Paulo, Brazil (http://www.esalq.usp.br/trilhas/palm/) and the sample of *E. precatoria* was collected at the Instituto Nacional de Pesquisas da Amazônia (INPA), in Manaus, AM, Brazil. The collections were registered according to the Brazilian laws (SISGEN number A411583, Brazil).

### Intact chloroplast isolation in sucrose gradient and chloroplast DNA extraction

The chloroplast organelles were isolated using the sucrose gradient method [36]. About 20 g of fresh leaves from each species were frozen with liquid nitrogen and macerated. The material was resuspended in 200 mL of isolation buffer (50 mM Tris-HCl pH 8.0, 0.35 M sucrose, 7 mM EDTA, 5 mM 2-mercaptoethanol and 0.1% BSA) and incubated for 10 min in the dark. The suspension was filtered through two layers of Miracloth (Merck), and the filtrate was centrifuged at 1,000 × g for 10 min.

The pellet was resuspended in 5 mL of isolation buffer and the suspension slowly laid out in the density gradients of 20/45% sucrose in 50 mM Tris-HCl (pH 8.0), 0.3 M sorbitol and 7 mM EDTA. The gradients were centrifuged at 2000 × g for 30 min, and the green band formed at the interface containing intact chloroplasts were collected. The solution containing the chloroplasts were diluted in three volumes of buffer and centrifuged at 3,000 × g for 10 minutes to obtain the purified chloroplasts in the pellet.

The pellet was resuspended in 2% CTAB buffer to promote lysis. The suspension was incubated and stirred at 65°C for 1 h. The supernatant was extracted twice with an equal volume of chloroform: isoamyl alcohol (24: 1) and centrifuged at 10,000 × g for 20 min. An equal volume of isopropanol was added and incubated at 20°C for 1h. Finally, the aqueous phase was centrifuged for 10,000 × g for 20 min, and the chloroplast DNA (cpDNA) pellet was washed with ethanol (70%), dried and resuspended with 40 μL TE (1 M Tris-HCl, 0.5 M EDTA, pH 8).

### Chloroplast genome sequencing, assembly and annotation

The genomic libraries were constructed using 100 ng of cpDNA and the Nextera DNA Flex kit (Illumina), following the manufacturer’s instructions. Paired-end sequencing (2x 150 bp) was performed on the Illumina NextSeq550 platform (Fundação Hemocentro de Ribeirão Preto, Brazil).

The assembly was conducted in three steps: First, the filtered reads from *Euterpe oleracea* were assembled in NOVOPlasty v 4.2 [37] (https://github.com/ndierckx/NOVOPlasty) using the *rbcL* gene sequence as a seed (NCBI accession number: MN621452.1) and the chloroplast genome of *Acrocomia aculeata* (NCBI accession number: NC_037084.1), a native Brazilian palm, as a reference to ordinate the contigs. Subsequently, all the three chloroplast genomes (*E. precatoria, E. edulis* and *E. oleracea*) were assembled in GetOrganelle v 1.7.3.1 [38] (https://github.com/Kinggerm/GetOrganelle/) using the *E. oleracea* chloroplast sequence obtained in NOVOPlasty as seed. Briefly, in GetOrganelle, the chloroplast genomes were assembled using default settings, starting with the recruitment of target-associated reads using Bowtie2 [39]. In this step, the seed chloroplast genome was used as target for the extension in five iterations. Then, the total target-associated reads were *de novo* assembled into a fasta assembly graph using SPAdes [40]. The *de novo* assemblies were trimmed and used to calculate all possible paths of a complete organelle genome [38]. Finally, the correctness and coverage of the assembly was assessed and confirmed in Geneious v2020 2.4. (https://www.geneious.com/, last assessed January, 2021). We used the “Map to reference” function to map the paired-end raw data onto the final assembled chloroplast genomes.

The chloroplast genome annotation was performed in GeSeq (Organellar Genome Annotation) [41] from the Chlorobox platform, with settings for the identification of protein coding sequence (CDS), rRNAs and tRNAs based on reference chloroplast sequences and homologies through BLAST search. Following the automatic annotation, a manual correction of start and stop codons and a verification of pseudogenes and intron positions were performed using GenomeView [42]. We then obtained the chloroplast circular genome maps using OGDRAW [43].

### Chloroplast genome structure comparison

To perform a comparative study and access the synteny between the obtained chloroplast sequences, we used a Perl script of MUMmer4 [44] (https://github.com/mummer4/mummer) to align the chloroplast genome from *E. edulis, E. oleracea* and *E. precatoria* with the function NUCmer. This analysis enables the identification of the conserved regions among sequences between species. The results were visualized in dot plots created by the function MUMmerplot. Additionally, to compare and align the *Euterpe* chloroplast sequences obtained with the GenBank sequences [34], MAFFT v.7 [45] was used as a way to identify differences between them. The location of the additional sequences found between them was also annotated in GeSeq [41].

Taking into account the species relationships from Arecoideae subfamily, and the previous evidence from palms chloroplast genome structures [28,46], we decided to conduct two multiple progressive sequence alignment in Mauve v.2.4.0 [47]. The first one only included chloroplast genomes from Brazilian native species of subfamily Arecoideae: *Euterpe edulis, Euterpe oleracea, Syagrus coronata, Astrocaryum aculeatum, Astrocaryum murumuru* and *Acrocomia aculeata*. The second analysis was carried out using 17 chloroplast genomes from different palm species (S1 Table) available in GenBank. Taking into the account the evolution of the group, we selected the species that represented the five palm subfamilies: *Phytelepas aequatoriallis* and *Pseudophoenix vinifera* (Subfamily: Ceroxyloideae); *Trachycarpus fortune* and *Caryota mitis* (Subfamily: Coryphoideae); *Nypa fruticans* (Subfamily: Nypoideae); *Calamus caryotoides* and *Eresmopatha macrocarpa* (Subfamily: Calamoideae); *Veitchia arecina* (Subfamily: Arecoideae) and the Brazilian native species also from the Arecoideae subfamily.

Expansions and contractions in the inverted repeats (IR) regions from the chloroplast genome structure were also explored. The chloroplast genome junctions (IRB/LSC; IRB/SSC, SSC/IRA; IRA/LSC) from the *Euterpe* species, and other Brazilian palm species from subfamily Arecoideae were examined to identify differences between individuals within the same genus and among subfamilies.

### Identification of SSRs, dispersed repeats, indels, SNPs and nucleotide divergence hotspots

Simple sequence repeats (SSR) consisting of 1-6 nucleotide units were carefully determined using the web package MISA (available at https://webblast.ipk-gatersleben.de/misa/) [48]. The criteria to search SSR motifs were: SSR of one to six nucleotides long, with a minimum repeat number of 10, 5, and 4 units for mono-, di-, and trinucleotide SSRs, respectively, and three units for tetra-, penta- and hexanucleotide SSRs. The SSRs sequences and location were compared among the species. The dispersed repeats (forward, reverse, palindrome and complement sequences) identified in REPuter [49] were based on the following criteria: minimum repetition size ≥ 30 bp and sequence identity ≥ 90% (Hamming distance = 3). Posteriorly, the position of the SSRs and repeats were manually compared with the gene annotation of each chloroplast genome.

MAFFT v.7 [45] was used to obtain pairwise alignments between the chloroplast genomes to pinpoint small insertions/deletions (indels) in the sequences. The alignment between the three species was also used to identify single nucleotide polymorphisms (SNPs) and nucleotide divergence hotspots with DnaSP v.5 [50]. Specific coding genes with a high number of SNPs were aligned with the software Muscle [51] in the Selecton Server [52], which was also used to identify the synonymous (Ka) or non-synonymous (Ks) mutations. With this analysis, according with the Ka/Ks ratios, it was possible to determine positive (Ka/Ks > 1) or purifying (Ka/Ks < 1) selection, estimated under the evolutionary model M8.

A sliding window analysis (window length of 200 bp and step size of 50 bp) was conducted to find nucleotide divergence hotspots. All the positions of indels, SNPs and divergence hotspots were manually identified using the annotation of the aligned chloroplast genomes, performed previously in GeSeq (Organellar Genome Annotation) [41].

### Phylogenomic studies

Plastid sequences of 54 palm species plus one outgroup (*Daypogon bromeliifolius*, Dasypogonaceae) (S1 Table) curated annotated features in Genbank format were separated into genes and entered into a SQLITE database using a custom Python script (available on demand from the corresponding author). Sequences of all putative coding proteins were then grouped by species and aligned with MAFFT, and later assembled into an interleaved NEXUS file for phylogenetic analyses. Species with missing sequences were filled with missing data in each gene matrix. Each block of gene sequences was then manually checked for start and stop codons and evidence of non-coding behaviour. For the coding genes, we annotated first, second and third positions for each codon, but the regions *cemA* and *rpl16* presented strange start codons and non-triplet insertions and were not annotated for codon positions. We tested three *ad hoc* partitioning schemes for models of molecular evolution: i) single model for the whole matrix, ii) four partitions: 1^st^, 2^nd^, 3^rd^ codon positions and a partition for *cemA* +*rpl16* (not split into codon positions), iii) five partitions: 1^st^, 2^nd^, 3^rd^ codon positions, *cemA* (not split into codon positions), *rpl16* (not split into codon positions). The assessment of molecular evolution model for each partition was calculated in the different partition schemes with the Akaike Information Criterion (AIC) in MrModelTest 2.0 [53], but they were all GTR+I+G, probably due to the complexity of mixing different genes in each partition. The assessment of the best partition scheme was then made using stepping stone (SS) [54] which is more accurate than taking the harmonic means of the likelihoods for model comparison. Bayesian inference were carried out with MrBAYES 3.2.7 [55] in the CIPRES platform [56], with one run of two chains, and 10 × 10^6^ generations, sampling one tree every 1,000. Marginal likelihoods of the SS runs were then compared using the standard Bayes Factors scale [57]. The same partition schemes were run separately to estimate the phylogeny with two runs of four chains (three hot and one cold), and 20 × 10^6^ generations, sampling one tree every 1,000 and discarding 25% initial generations for burn-in. The remaining trees were checked for estimated sample sized (ESS) > 200 in all parameters, and the final majority-rule was computed with MrBayes, rooted with the single outgroup *D. bromeliifolius* and the tree was exported to FigTree 1.4 [58] for drawing, and later improved with InkScape [59] for designing the figures.

### Genetic variation and comparative analyses with nuclear and mitochondrial sequences

After the assembly of the chloroplast genomes, the reads were also used to obtain SNPs for preliminary comparative analysis between species. Initially, the raw reads of each species were aligned in their respective assembled chloroplast genome using Bowtie2 [39]. Subsequently, the non-aligned sequences (non-chloroplast) were obtained with SAMTools [60], and converted to fastq file using Picard Tools program [61].

The reads corresponding to non-chloroplast genome (nuclear and mitochondrial data) were cleaned using *process_shortreads* from the Stacks package v.1.42 [62]. Sequences from each species were used to build loci with a minimum depth of three and up to two mismatches, applying the *ustacks* program (-m 3, -M 2) [62]. Subsequently, a catalog was built from each species, allowing for up to two mismatches (-n 2, cstacks) [62], and using *sstacks* [62] loci were matched with the catalog of each species. *Rxstacks* [62] was used as a correction step, assuming a mean log likelihood of -10 to discard loci with lower probabilities. The *populations* [62] program was administered for a final filtering to retain loci with a maximum missing data of 5%, and also used to verify the number of private alleles (*Ap*), observed heterozygosity (*H*_*O*_) and expected heterozygosity (*H*_*E*_; S1 Fig).

Finally, the mean sequence depth and the mutation counts of the SNPs were determined in vcftools [63]. The number of alleles (*A*) in each species and the genetic differentiation (*F*_*ST*_) between them were estimated with *Adegenet* v 1.3 -1 [64] and *Genpop* v 1.1.7 [65] for the platform R v 4.0.3 [66].

## Results

### Organization of the three *Euterpe* species chloroplast genomes

The chloroplast genomes of *E. edulis, E. oleracea* and *E. precatoria* had the typical quadripartite structure (Fig 1), with the presence of two copies of inverted repeat regions (Inverted Repeats A and B = IRA and IRB) and two single copy regions (Large Single Copy = LSC and Small Single Copy = SSC). *E. precatoria* had the largest chloroplast genome size (159,275 bp) and the largest LSC and SSC (87,282 bp and 17,756 bp, respectively), while *E. edulis* had the largest IR regions and *E. oleracea* a slightly larger amount of GC content (37,3%) (Table1). All three chloroplast genome annotations resulted in the identification of 113 unique genes, with 30 tRNAs, 4 rRNAs and 79 protein coding genes (Fig 1, Table 2).

**Fig 1.**
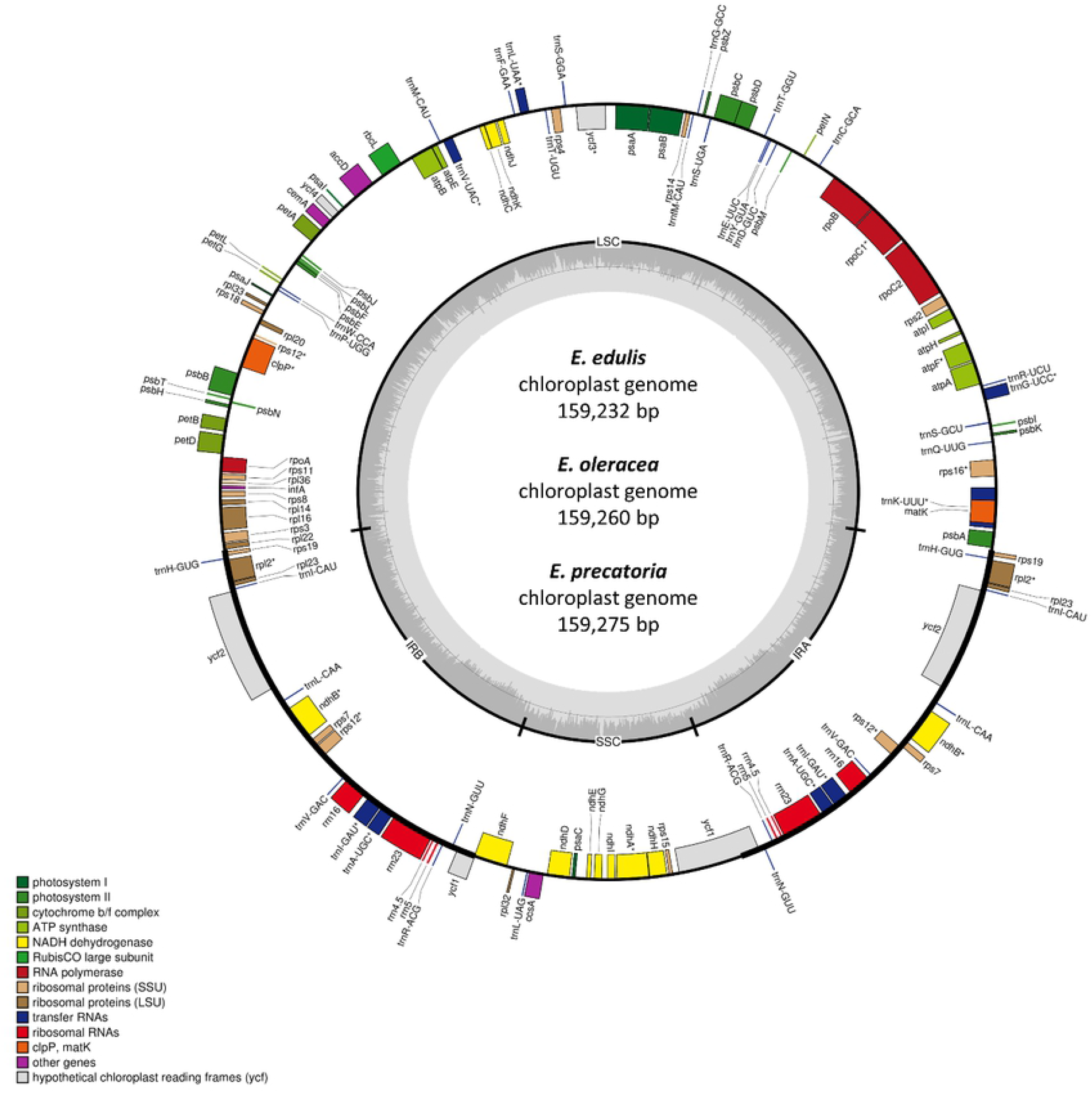
Gene map of the *Euterpe edulis, E. oleracea* and *E. precatoria* chloroplast genomes. Genes represented inside the large circle are oriented clockwise and the ones outside are oriented counter clockwise. The different colors represent functional groups, and the darker gray in the inner circle indicates the GC content. The quadripartite structure is also reported as: LSC = Large Single Copy, SSC = Small Single Copy, IRA/IRB = Inverted Repeats A and B.

Considering the duplicated genes in the IRs, more than 50% of the three chloroplast genome sequences are from protein coding regions (86 genes, Table 1 and Table 2). Among genes, *ycf2* presented a slight difference between species, 6,903 bp in *Euterpe edulis* and 6,879 bp in *E. oleracea* and *E. precatoria*. Furthermore, chloroplast tRNA and rRNA were conserved among species, constituting 1.8% and 5.7% of their sequences, respectively. Introns constituted ca. 12% and intergenic regions represented almost 31% of the chloroplast genomes (Table 1).

**Table 1.**
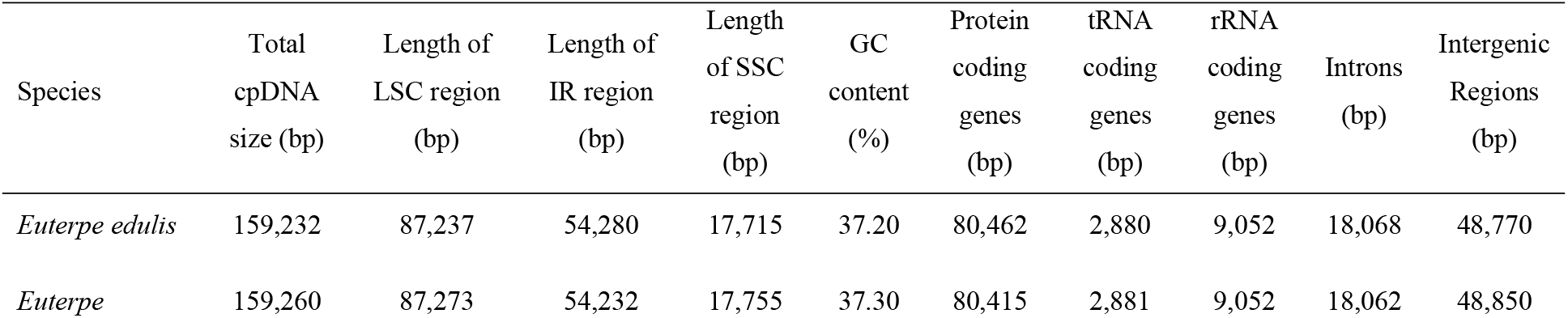
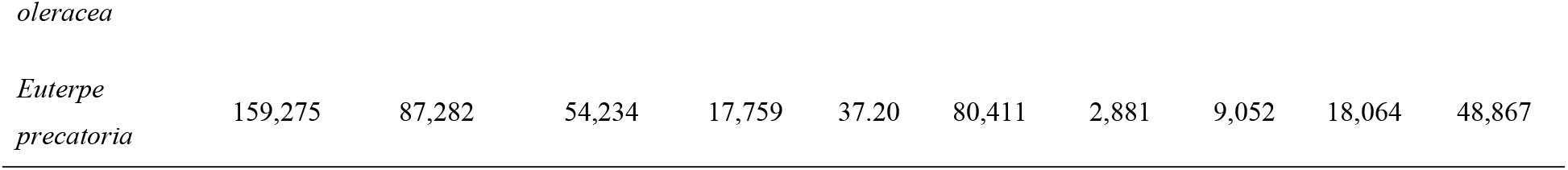
General features of chloroplast genomes of three *Euterpe* species.

**Table 2.**
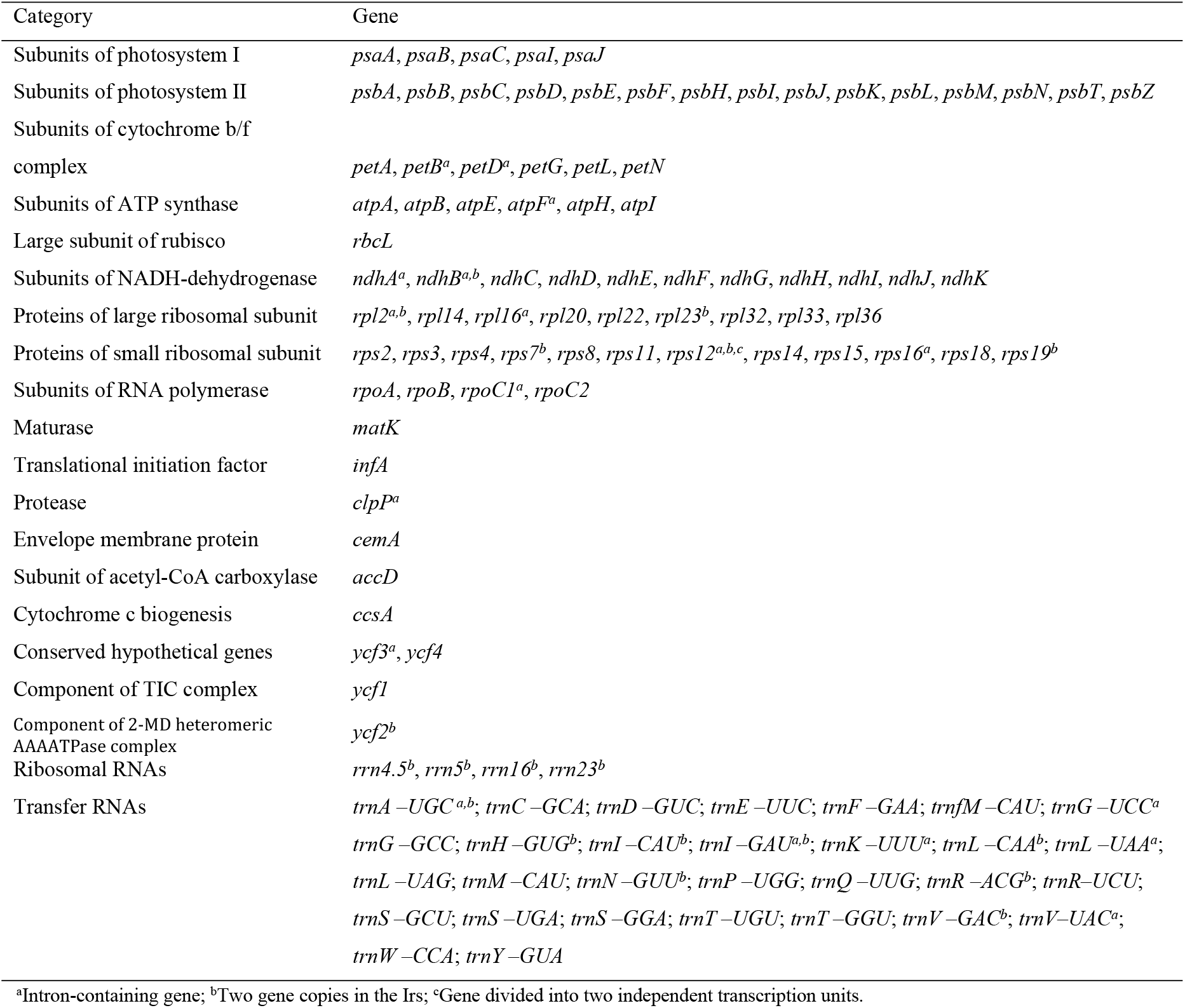
Gene content in *Euterpe edulis, E. oleracea* and *E. precatoria* chloroplast genomes according to each respective category.

All chloroplast genomes had 17 unique genes (11 protein- and six tRNA-coding genes) with introns, five duplicated genes in the inverted repeats regions with introns, and two introns in the *ycf3* and *clpP* genes. Accounting for the lengths, the largest intron was identified in *trnK – UUU* (2,621 bp *– E. edulis*; 2,615 bp – *E. oleracea*; 2,618 bp – *E. precatoria*) and the smallest in *trnL-UAA* (519 bp – *E. edulis*, 519 bp – *E. oleracea*, 521 bp – *E. precatoria*).

### Chloroplast genome structures and comparative analyses

The comparative analysis enabled us to identify a high level of synteny between the three *Euterpe* chloroplast genomes, with large conserved blocks. The only structural difference among *Euterpe* chloroplast genomes were three small inversions and single nucleotide polymorphisms (SNPs) between *E. edulis* and *E. oleracea* chloroplast sequences (S2A, B and C Figs). In a local alignment we detected these divergences in the region between 66,081 to 69,440 bp; the first corresponding to an intergenic region between *petA* and *psbJ* genes from *E. oleracea*; the second found in the *psbJ* gene and the third in the intergenic region of *trnW-CCA* and *trnP-UGG* from *E. oleracea* (S2D Fig). The parallel between the chloroplast genomes of *Euterpe edulis* and *Euterpe oleracea* with the ones available in Genbank [34] showed differences in the total number of base pairs. *E. edulis* and *E. oleracea* were 835 bp and 458 bp, respectively, larger than the others. The alignment revealed that these increases were due to insertions that occurred mainly in intergenic spacers (83.3% of insertions in *E. edulis* and 100% in *E. oleracea*; S2 Table).

Although exhibiting high synteny and only a few structural rearrangements in their chloroplast genome, the most notable divergence among the three Brazilian native species from subfamily Arecoideae was in the length of the LSC, between 40,000 and 50,000 bp (Fig 2). A high level of synteny was also observed in a comparison of palm chloroplast genomes from 12 genera, with five subfamilies (S3 Fig). *Astrocaryum aculeatum, A. murumuru* and *Syagrus coronata* are native palms from subfamily Arecoideae with smaller LSCs compared with *Euterpe* palms (Fig 2 and S3 Fig). *Acrocomia aculeata*, also a Brazilian palm, presented a reduction in the size of the region comprising the genes *ndhJ, ndhK, ndhC, trnF-GAA* and *trnL-UAA* (in lime green Fig 2). Additionally, *Astrocaryum aculeatum* and *A. murumuru* showed a flip-flop recombination in this same region (in lime green, Fig 2) [67].

**Fig 2.**
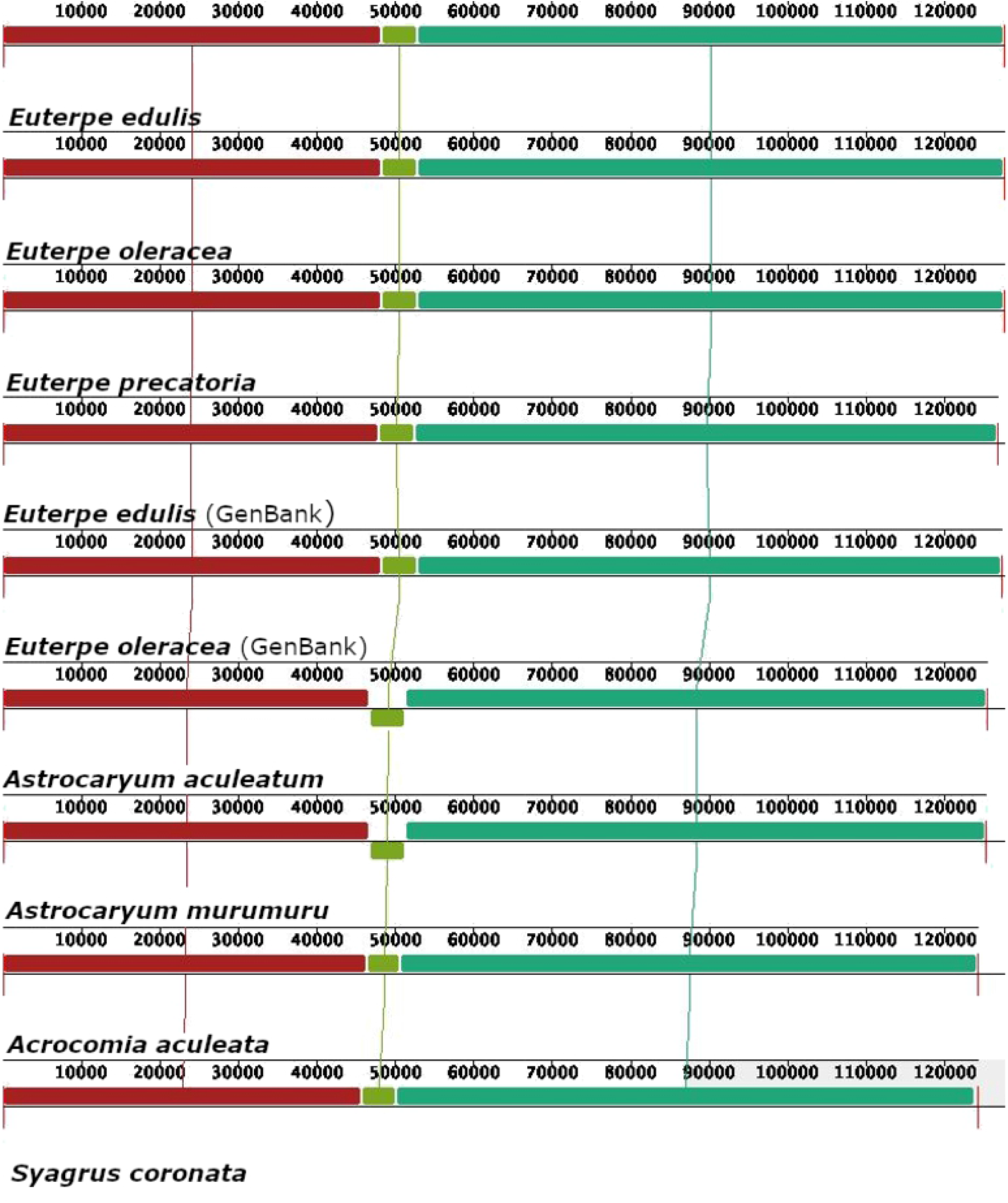
Synteny and divergence in the SSC size detected in Arecaceae chloroplast genomes using the Mauve multiple-genome alignment program. A sample of nine different chloroplast genomes is shown. Color bars indicate syntenic blocks and the lines indicate the correspondence between them. Blocks on the top row are in the same orientation, while blocks on the bottom row are in inverse orientation.

In addition, we explored in more details the chloroplast IR expansions and contractions of the Brazilian palm species from the subfamily Arecoideae (Fig 3). Comparing the IR borders from Genbank available for *Euterpe edulis* chloroplast genome, it was possible to observe a small difference in the *rpl22*-*rps19* location and an increase in the length of *rps19*-*psbA* intergenic spacers. Among the chloroplast genomes from *Euterpe* genus, we identified an expansion in the *ndhF* gene at the IRB region from *Euterpe precatoria*. Among the Brazilian palm species, a contraction in the IR from *Syagrus coronata* influenced the position of *rps19*. Among all species analyzed, the most common variations were identified around the positions of *rpl22-rps19* (IRB-SSC), the *ycf1* length (IRB and IRA), and the *rps19-ycf1* (IRA-LSC) (Fig 3).

**Fig 3.**
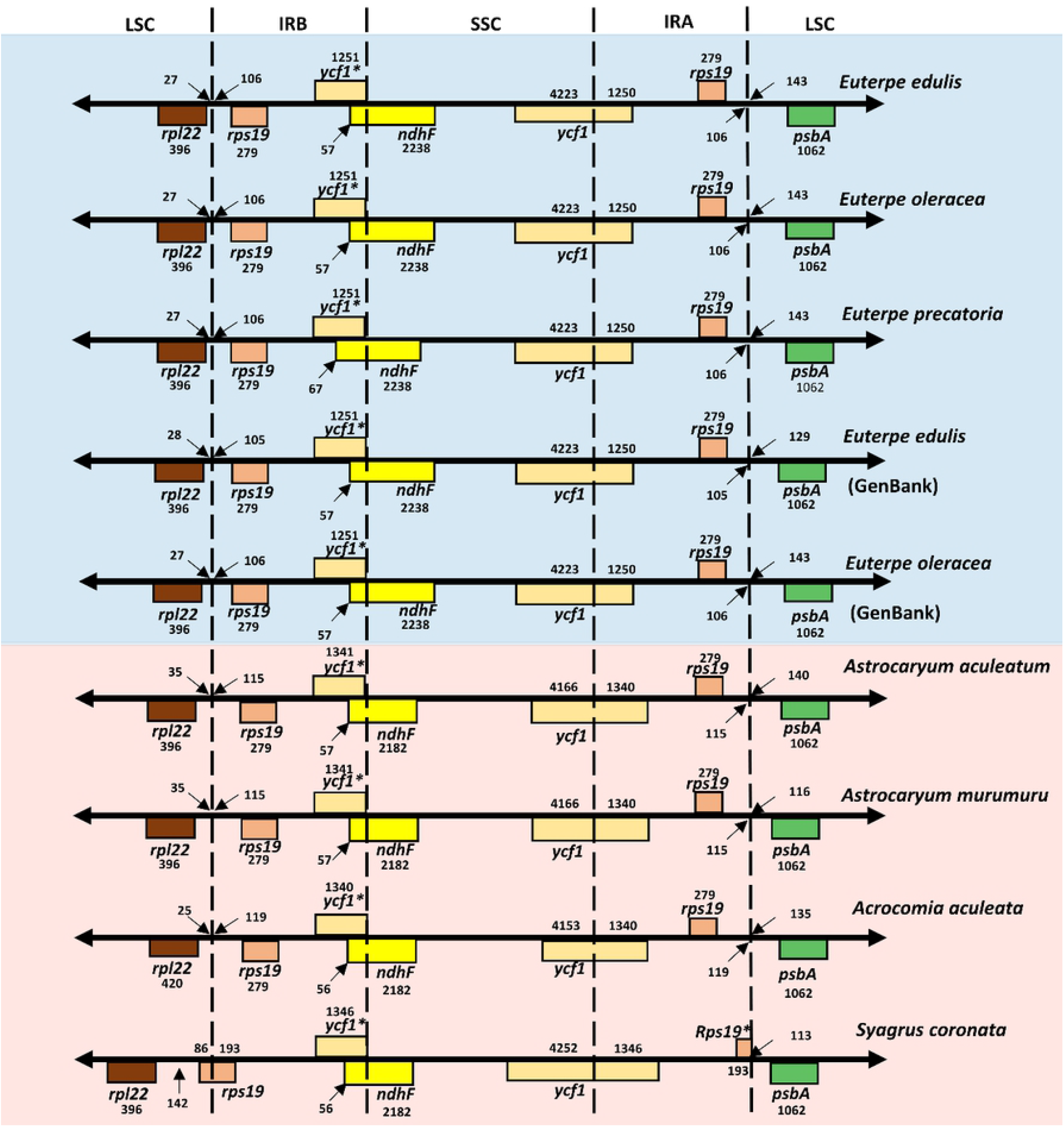
Comparison of the IRA and IRB borders among Brazilian palms from the Arecoideae species. The numbers indicate the lengths of IGSs, genes, and spacers between IR-LSC and IR-SSC junctions. The *ycf1\** and *rps19\** genes have incomplete CDSs.

### Sequence repeats and polymorphisms between *Euterpe* chloroplast genomes

We identified a total of 323 SSRs (*E. edulis*: 111, *E. oleracea*: 105, *E. precatoria*: 107 SSRs) in the chloroplast genomes from the three species. Most of them were located in intergenic spacer regions (IGS: 72.07% *E. edulis*, 72.38% *E. oleracea*, 71.96 % *E. precatoria*; Fig 4A), especially in the IGS between the tRNAs trnS-GCU/trnG-UCC (S3 Table). The SSRs were more abundant in the LSC region of the chloroplast genomes (78.38% *E. edulis*, 78.10% *E. oleracea*, 77.57% *E. precatoria*; S3 Table) and less frequent in the IR regions. They were mostly mononucleotides (62.16% *E. edulis*, 62.86% *E. oleracea*, 60.75% *E. precatoria*; Fig 4B) and composed with A/T motifs (60.36% *E. edulis*, 60.00% *E. oleracea*, 59.81% *E. precatoria*; S4A Fig). From the 323 SSRs, 47 had the same sequence and gene location among *E. edulis, E. oleracea* and *E. precatoria*. We also identified 62 SSRs between *E. edulis/E. oleracea* and *E. edulis/E. precatoria*, and 68 between *E. oleracea and E. precatoria* (S4 Table).

**Fig 4.**
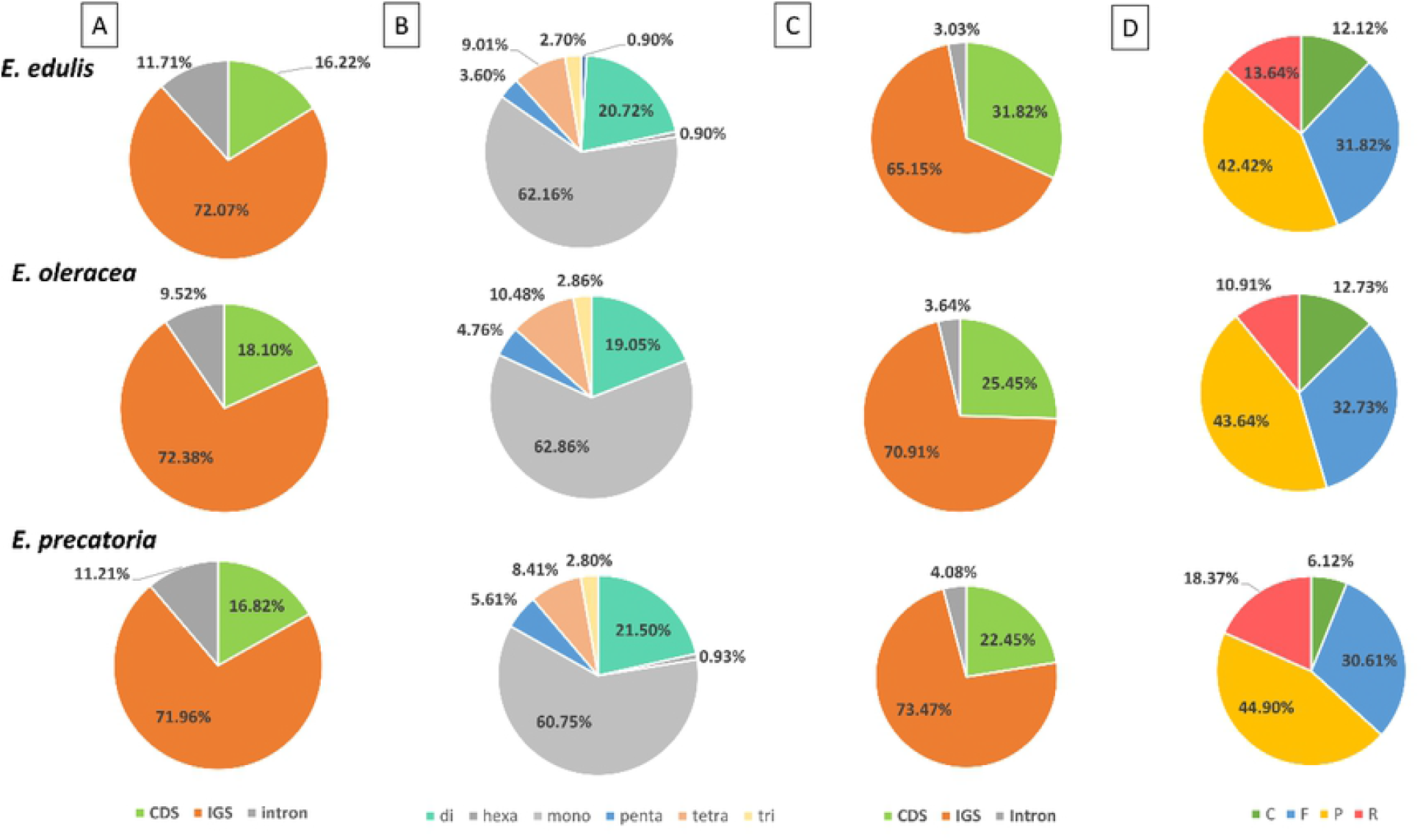
Distribution and classification of SSR and dispersed repeats in the chloroplast genomes of *Euterpe edulis, E. oleracea* and *E. precatoria*. (A) Proportion of coding and non-coding regions containing SSRs; (B) Proportion of different types of SSR present in the chloroplast genomes; (C) Proportion of regions containing repeats; (D) Frequency distribution of different types of repeats: F = Forward, P = Palindrome, R = Reverse and C = Complement. CDS = Coding sequence, IGS = Intergenic spacer.

Considering the dispersed repeats, we observed 66 in *E. edulis*, 55 in *E. oleracea* and 49 in *E. precatoria*, ranging from 30 to 77 bp (S4B Fig). These repeats were distributed mostly in the IGS regions (65.15% *E. edulis*, 70.91% *E. oleracea*, 73.47% *E. precatoria*; Fig 4C). Comparing the species, in *E. precatoria*, most of the repeats were found in the intergenic region of *psaC*/*ndhE*. However, in *E. edulis* and *E. oleracea* the repeats were concentrated in the *ycf2* gene (S5 Table). Regarding repeat types, most of the repeats are palindrome or forward sequences, 40% and 30%, respectively (Fig 4D). They were also mainly found in the LSC region (48.48% *E. edulis*, 52.73% *E. oleracea*, 44.90% *E. precatoria*; S5 Table).

Based on the pairwise alignment between the species chloroplast genomes, we identified 93 indels between *E. edulis*/*E. oleracea*, 103 between *E. edulis/E. precatoria* and 58 between *E. oleracea*/*E. precatoria*. Most indels were distributed in the IGS (Fig 5A), especially between the *E. oleracea/E. precatoria* chloroplast genome, located mainly in the LSC region (80.65% ∼ 72.41%; S6 Table). The highest occurrence of indels was identified in the *trnS-GCU*/*trnG-UCC* of the three alignments (S6 Table).

**Fig 5.**
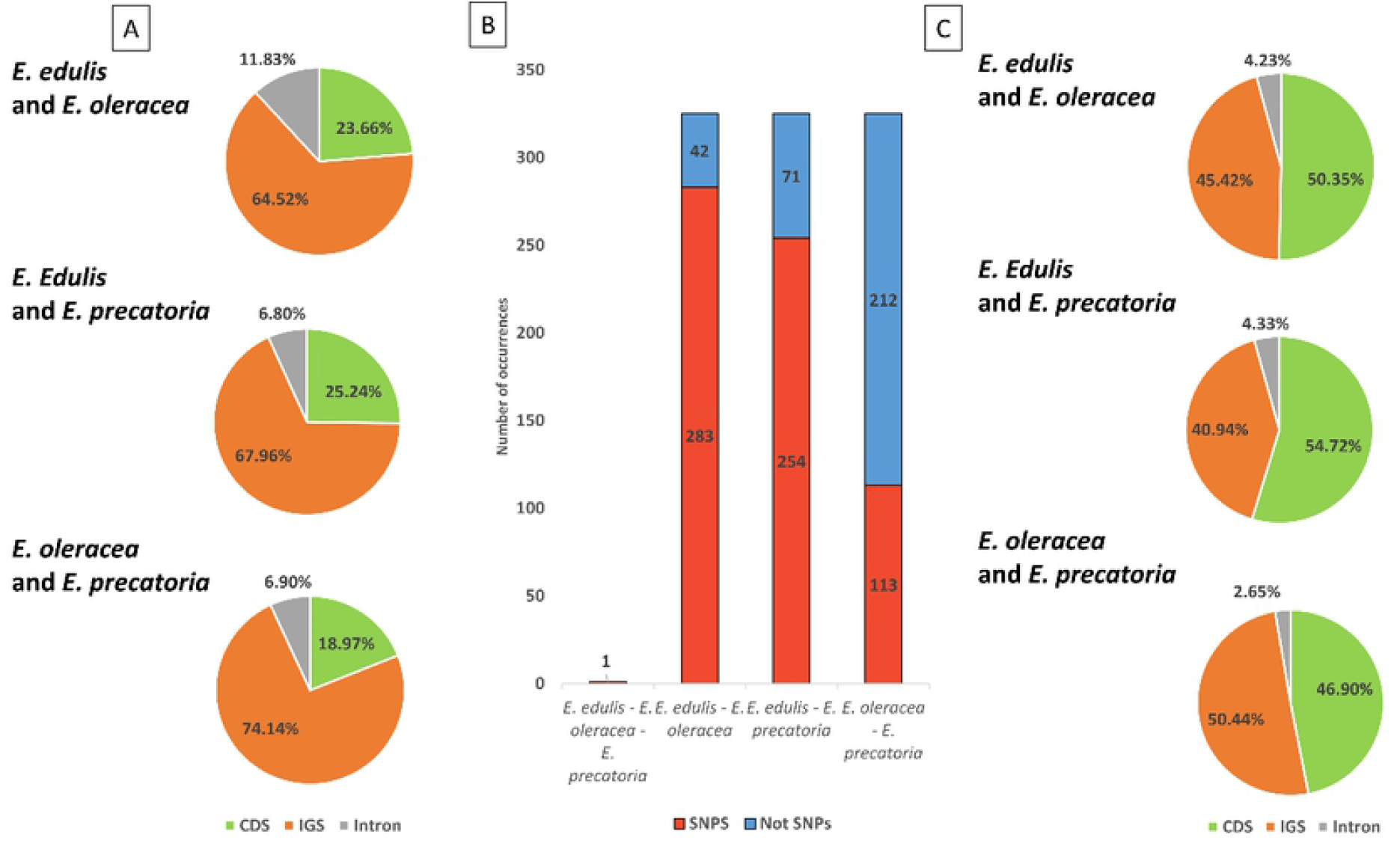
Indels and single nucleotide polymorphisms (SNPs) detected in comparisons between *Euterpe* chloroplast genomes. (A) Proportion of indels in different coding and non-coding regions; (B) Comparison of the number of SNPs found in the alignment (C) Proportion of SNPs in different coding and non-coding regions of the chloroplast genomes.

A total of 325 SNPs were detected between the three species (Fig 5B). In a pairwise comparison, 283 SNPs were detected between *E. edulis/E. oleracea* and 254 SNPs between *E. edulis/E. precatoria*, which were most frequent in the CDS regions (50.35% and 54.72%, respectively; Fig 5C). Between *E. oleracea/E. precatoria*, a lower number of SNPs was found (113 SNPs), having the greatest number concentrated in the IGS (50.44%; Fig 5C). Only one SNP was found shared among the three species (Fig 5B).

The large number of SNPs in the CDS region occurs mainly in the *atpE, ycf1* and *psbJ* genes (S7 Table). These *atpE, psbJ* and *ycf1* genes presented respectively 19, 18 and 18 SNPs for *E. edulis/E. oleracea* and 20, 19 and 21 SNPs for *E. edulis/E. precatoria*. For *E. oleracea/E. precatoria* there were only 1 (*atpE*) and 7 (*ycf1*) SNPs (S7 Table).

The LSC region had the highest number of SNPs in the chloroplast genomes (69.01% *E. edulis/E. oleracea*, 66.54% *E. edulis/E. oleracea*, 64.91% *E. oleracea/E. precatoria*; S7 Table). For *E. edulis/E. oleracea* the greatest number of SNPs corresponded to C/T - T/C (74) substitutions, the same was saw for *E. edulis/E. precatoria* T/C - C/T (66), and 29 substitutions of G/A - A/G and C/T - T/C was observed for *E. oleracea/E. precatoria* (S5 Fig).

Considering pi > 0.02, the sliding window analysis revealed six hotspots of high nucleotide polymorphism among the *E. edulis, E. oleracea* and *E. precatoria* chloroplast genome sequence. The six hotspots were in IGS regions (Fig 6). Among them, the two hotspots with the highest polymorphism were located between the *trnM*/*atpE* (Pi > 0.06; Fig 6) and *psbJ*/*psbL* (Pi > 0.06; Fig 6). Comparing with the SNP identification, the *atpE* gene had the highest number of SNPs among the species *E. edulis/E. oleracea* and *E. edulis/E. precatoria*. The *psbJ* gene had 18 SNPs in *E. edulis/E. oleracea* and 19 SNPs in *E. edulis/E. precatoria*.

**Fig 6.**
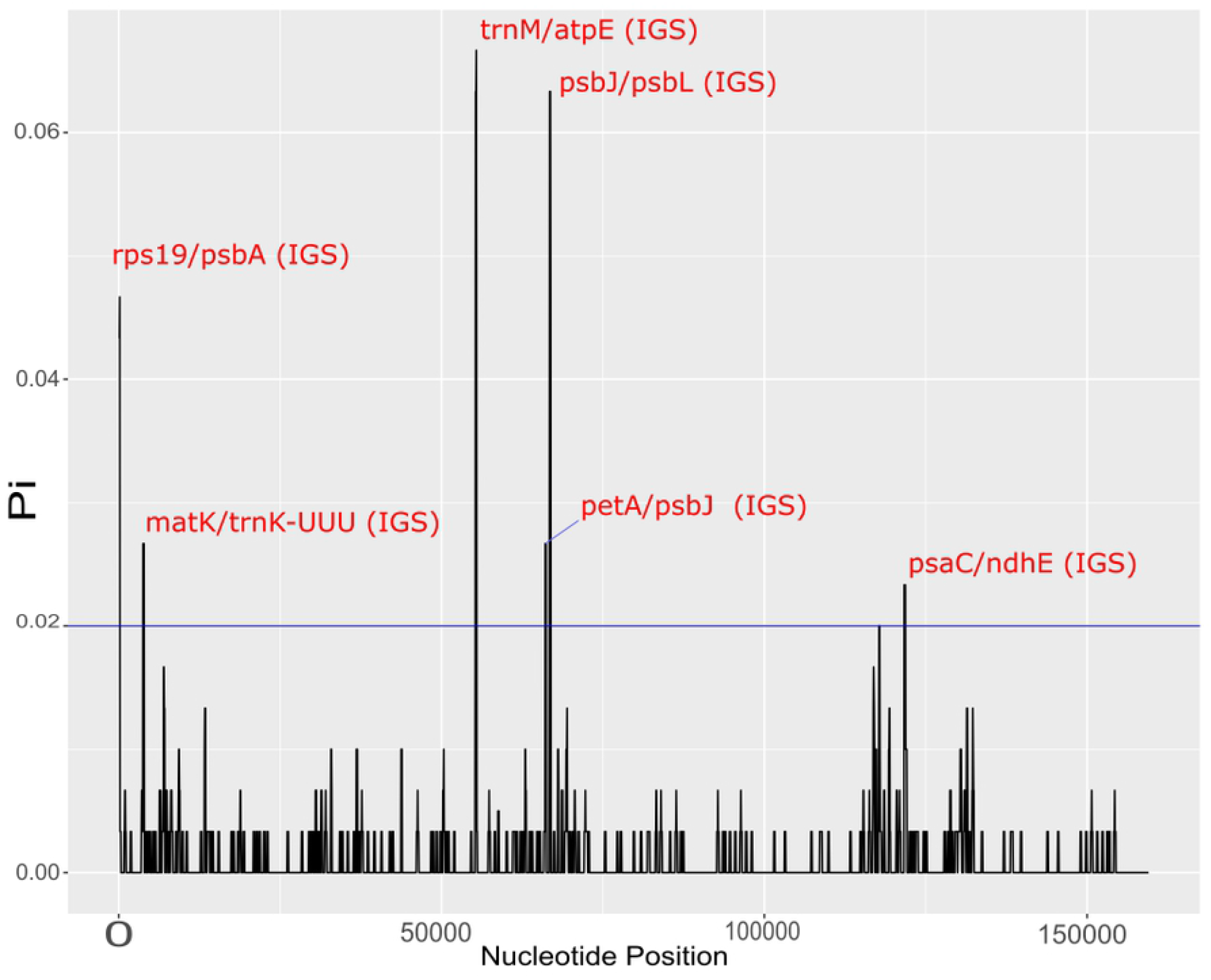
Sliding window analysis of the alignment from the chloroplast genome of *Euterpe edulis, E. oleracea* and *E. precatoria* chloroplast genomes. The regions with high nucleotide variability (Pi > 0.02) are indicated. Pi is the nucleotide diversity of each window, and the window length was 200 bp with 50 bp step sizes.

Both genes, *atpE* and *psbJ*, were analyzed for positive or purifying selection across the palm phylogeny using the estimation of Ka/Ks ratio. In both cases, the Ka/Ks< 1 revealed that no positively selected sites were found in the corresponding protein sequences, indicating purifying selection. The analysis of the amino acid sequence resulted in Ka/Ks rates varying from 0.17 to 0.87 for *atpE* (S8 Table) and ranging from 0.50 to 0.96 for *psbJ*. Also, from the 134 amino acids of the *atpE* protein, 68 had Ka/Ks < 0.19 and from the 40 amino acids, 37 had Ka/Ks < 0.55, in the case of *psbJ* (S9 Table).

### Phylogenomic of *Euterpe* based on chloroplast sequences

In the phylogenomic analysis, the marginal mean likelihoods of partition schemes for the two runs were i) -186887.39, ii) -178507.00, and iii) -178497.59. Therefore, the best partition schemes were by far the ones considering codon positions, although ‘iii’ was strongly better than ‘ii’ with the 2*log difference = 18,82 (>10 is strong in Kass and Raftery 1995). We also compared the majority-rule consensus trees of the three partition scheme analyses, which did not differ in topology, but in general the scheme ‘iii’ presented better support in some nodes. For that reason, the figure presented and used in our discussion will be based on partition scheme ‘iii’.

The phylogeny using coding sequences from 54 whole chloroplast genomes from palm species rooted in the outgroup species *Dasypogon bromeliifolius* revealed the same relationships in subfamilies previously reported [34], with highly supported nodes (Fig 7). According to our results, the species from *Euterpe* were placed in subfamily Arecoideae, tribe Euterpeae, and were sister to tribe Areceae. The new samples of *E. oleracea* and *E. edulis* were grouped with the respective previously published chloroplast genomes and the *E. precatoria* was placed in a branch sister to *E. oleracea*.

**Fig 7.**
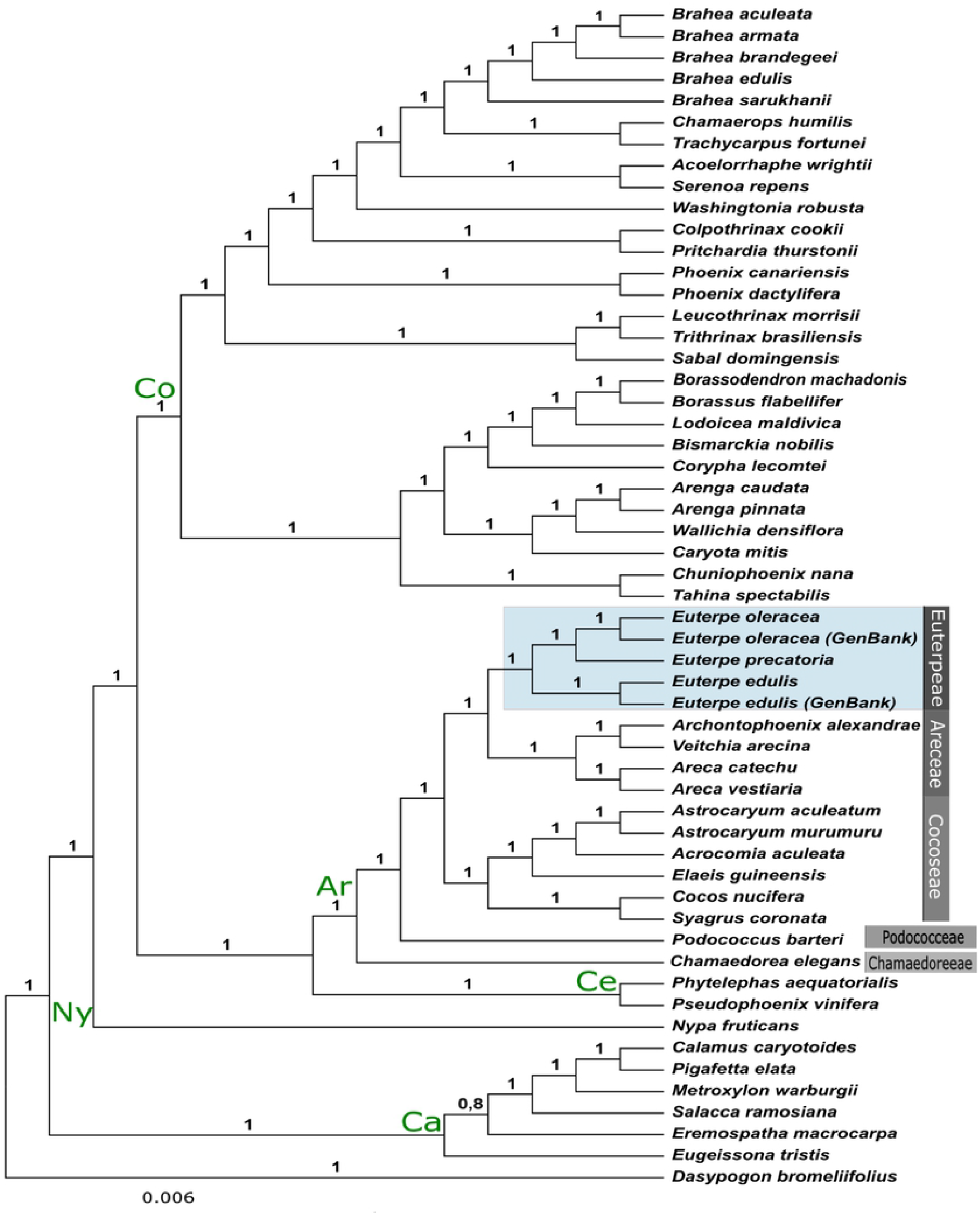
Majority-rule consensus tree of 30,000 trees obtained from a Bayesian inference analysis of chloroplast protein coding genes of 55 taxa. Posterior probabilities (PP) for each are indicated above branches. Co = Coryphoideae, Ar = Arecoideae, Ny = Nypoideae, Ce = Ceroxyloideae, Ca = Calamoideae.

### Species divergence with nuclear and mitochondrial genome

From the 11,738,792 reads that were generated for *E. edulis*, 553,857 (4.71%) aligned to the assembled chloroplast genome. The remaining (ca. 95%) reads correspond to nuclear and mitochondrial regions. Similar proportions were also found for *E. oleracea* and *E. precatoria* for which 12,987,827 and 12,612,001 reads were generated, respectively. Moreover, 249,387 (1.92%) and 132,227 (1.04%) were aligned to the assembled chloroplast sequence for both species, respectively. Using Stacks, we found 1,077 SNPs in the nuclear and mitochondrial genome sequences, with 10X mean sequencing depth per locus (S6A Fig). Among *Euterpe* species, more transitions than transversions were detected, and the most frequent mutations were A-G and C-T transitions (S6C Fig).

The highest number of alleles (*A* = 1,309) and highest observed heterozygosity (*H*_*O*_ = 0.162) was identified in *E. oleracea*, while the highest number of private alleles was observed in *E. precatoria* (*Ap* = 421). All the three species had a higher value of *H*_*O*_ compared with the expected heterozygosity (*H*_*E*_, Table 3).

**Table 3.** Parameters of genetic diversity using 1,077 SNPs found the nuclear and mitochondrial genome sequences from three *Euterpe* species.

|  | *E. edulis* | *E. oleracea* | *E. precatoria* |
| --- | --- | --- | --- |
| $A$ | 1,251 | 1,309 | 1,263 |
| $Ap$ | 292 | 260 | 421 |
| $H_O$ | 0.162 ($\pm 0.011$) | 0.215 ($\pm 0.013$) | 0.173 ($\pm 0.012$) |
| $H_E$ | 0.081 ($\pm 0.006$) | 0.108 ($\pm 0.006$) | 0.086 ($\pm 0.006$) |

$A$ = number of alleles, $Ap$ = number of private alleles, $H_O$ = observed heterozygosity, $H_E$ = expected heterozygosity.

The pairwise *F*_*ST*_ showed high and significant divergence between species, with values higher than 0.770. Although all the *F*_*ST*_ values were high, the species *E. edulis* and *E. precatoria* presented the greatest genetic divergence (0.860, S7 Fig) among the three pairs of species, and *E. edulis* and *E. oleracea* had the lowest divergence (0.779, S7 Fig).

## Discussion

### High level of conservation between Brazilian palms chloroplast genomes

Among the chloroplast genomes from the *Euterpe* genus, we found a conserved organization and gene content, with small differences between species, as in the length of the duplicated *ycf2* gene. The *ycf2* is the largest plastid gene reported in angiosperms [68], and is part of the 2-MD heteromeric AAA-ATPase complex directly interacting with various translocating preproteins [69]. Particularly, *ycf2* has an insertion in the *E. edulis* chloroplast genome that is absent in *E. oleracea* and *E. precatoria*, a feature previously observed in *Euterpe* chloroplast genomes [34]. Across palm species, the chloroplast genome structure is highly conserved as reported in other studies [28,34,70]. The multiple alignments with 15 different palms from five subfamilies did not exhibit significant rearrangements in the chloroplast structure. The only rearrangement observed was an inversion of 4.6 kb in the *Astrocaryum* chloroplast genome. It was possibly to have a lineage-specific structural variation, creating a isoform from a flip-flop recombination between inverted repeats [67]. However, differences in the total size of chloroplast genomes were observed in the same species, such as in *E. edulis* and *E. oleracea*, which may be due to the assembly method used. Different from the approach of all possible paths’ calculation to complete target organelle genome, as applied by GetOrganelle, other widely used pipelines utilize reference genomes to select and filter scaffolds/contigs for further concatenation or post-assembly gap filling and closing [38].

Besides these conserved general features, similar occurrences of SSRs were identified among the three *Euterpe* species. Currently, SSR markers are very useful in studies of population structure, genetic mapping, and evolutionary processes [71]. The SSR identified here can be a valuable resource for studies aiming at the conservation of the species or their sustainable exploitation. These SSR are mostly composed of A/T motifs, and are more frequently located in the IGS of the single-copy regions from the chloroplast genomes [1,72,73], which is expected since these regions evolve faster than the CDS [74]. Also, the limited occurrence of SSRs in the IRs is related to a lower mutation rate observed in these regions, caused by an efficient mechanism of gene copy-correction, as observed in other palm chloroplast genomes [34,75].

The dispersed repeats (forward, palindrome, reverse and complement) showed the same pattern of the SSRs and were frequently distributed in the IGS of the single copy regions, differing from a predominant occurrence of reverse repeats (18,37%) in *E. precatoria*. Actually, the number of repeats found in the *Euterpe* chloroplast genomes were not considered high [76]. According to Milligan et al. [77], repetitive sequences are substrates for recombination and chloroplast genome rearrangements. Since non-structural rearrangements were detected in the *Euterpe* chloroplast genomes, low frequency of repetitive elements was also expected.

Regarding the structure patterns of the IRs, we could identify most of the variation related to expansion/contractions when comparing only Brazilian palm species from subfamily Arecoideae. Events of expansion and contraction at the IR boundaries was previously observed in the *ycf1* gene at IR-SSC/LSC junction and *rps19* intergenic spacer with *rpl22*/*psbA* in IR-SSC/LSC junctions of other species [28,75].

### Indels and single nucleotide polymorphisms clearly demonstrate differentiation among the three *Euterpe* species, which was reflected in the phylogeny

We identified six hotspots of high nucleotide polymorphism with the largest polymorphism (Pi > 0.06) in intergenic regions composed mainly by genes with a great number of SNPs. The genes *atpE* and *psbJ* had the greatest number of SNPs, especially when considering the nucleotide substitutions between *E. edulis* and the *Euterpe* Amazonian species, *E. oleracea* and *E. precatoria*. This divergence between *E. edulis* in relation to the two Amazonian species was also observed in the number of indels and plastid SNPs and was almost 50% higher than the number of polymorphisms detected in a pairwise analysis with *E. oleracea* and *E. precatoria*.

Indels display no ambiguity in complex mutation patterns and SNPs are the most abundant type of markers [78], giving us a clear pattern of divergence between *E. edulis* in relation to *E. oleracea* and *E. precatoria*. In our study, these mutations were frequently found in the CDS than in the intergenic spacers. This high number of indels and SNPs between *E. edulis/E. oleracea* had already been identified in a previous study [34], but they were mainly found in non-coding regions, which presumably may be related to a more recent event of divergence.

Regarding the phylogenetic relationships among the palm species, we identified a clear distinction among subfamilies using the informative chloroplast coding sequences, corroborating previous studies [28,34,67,79]. Our phylogeny suggests that *E. oleracea* and *E. precatoria* are sister species with an early split of *E. edulis* from these two, differently from what was observed previously, having four nuclear low copy markers and one plastid region [33]. Our result reveals the possibility of incongruences between nuclear and plastid phylogenies in *Euterpe*, since the previous study using nuclear markers reported paraphyly in *E. precatoria* and showed one of the *E. precatoria* varieties as sister to *E. edulis* [33].

The closest relationship between *E. oleracea* and *E. precatoria* could reflect their environmental occupation since both species occur in the Brazilian Amazon. *E. edulis*, nonetheless, evolved via vicariant split from its common ancestor (around 1.4 Mya), and the speciation was intensified by the savanna barrier composed from the Cerrado and Caatinga biomes [33,34,80]. Furthermore, the phenotypic plasticity and local adaptation may have favored the successful expansion of *E. edulis* throughout the Atlantic Forest [11,80].

Regarding the coding sequences, the *rpl22, rpl32, rpl14, rps14, matK, accD, rbcl, ccsA, ycf2 and ycf1* genes have been reported to harbor positive selection in an analysis using 41 species of the Arecaceae family, including the *E. oleracea* and *E. edulis* chloroplast genomes [34]. This was highlighted as indicative of convergent evolution, associated with environmental adaptation in *E. edulis*. However, using 52 palm species, we could only identify a tendency of purifying selection for *psbJ* and *atpE*, the genes with higher SNPs and polymorphisms hotspots. This may reflect the typically conservative nature of the chloroplast genome sequences across most angiosperms [81,82]. These genes are essential for plant development, since *atpE* encodes a subunit of the ATP synthase complex that participates in the photosynthetic phosphorylation [83] and *psbJ* is associated with photosystem II (PSII) [84].

### A comparison using nuclear and mitochondrial genome sequences detected high divergence between *Euterpe* species

The reads from nuclear and mitochondrial DNA obtained from the sequencing were used in this first-time comparison of the genetic diversity and genomic divergence among the three species from the *Euterpe* genus. The advances in next-generation sequencing (NGS) is allowing the discovery of a large number of single nucleotide polymorphism (SNPs), even for non-model species [85–88]. This generates a valuable source of bi-allelic genetic markers that are appropriate to evaluate the genetic diversity even when sample sizes are small [89]. Additionally, SNPs occurs at higher density across the genome, with lower genotyping error rates than other markers [90], enabling robust estimates of genetic diversity and structuring [9].

The three *Euterpe* species had similar number of alleles and low observed heterozygosity. Yet, they are higher than the expected heterozygosity, with *E. oleracea* presenting slightly higher results. However, despite this enlightening information obtained with the remaining nuclear and mitochondrial sequences, we highlight that these findings could be biased because they were based on only one individual per species. Considering assessments of genome-wide diversity, previous studies identifying SNPs in palms were only performed for *E. edulis* [9,80]. Furthermore, the genetic diversity estimates from our *E. edulis* sample, that was also from the Atlantic rainforest of São Paulo state, were in agreement with previously published studies. This observed genetic variability could be influenced by the type of vegetation and the altitude where the samples were collected [9,80].

In summary, our results open the way for future genome-skimming studies in palms. Also, the addition of larger sample sizes from different sampling locations will provide a better understanding on the evolution and diversification of this important group of plants. However, the high differentiation found between the three *Euterpe* species may indicate that they are evolving independently and that this differentiation is caused by a private pool of alleles from each species, which is in accordance with the phylogenetic analyses.

## Acknowledgements

The authors would like to thank Dr. Charles R. Clement for sending samples of *Euterpe precatoria* from Instituto Nacional de Pesquisas da Amazônia (INPA), Manaus, Brazil for the genomic extraction and for reading the manuscript.

## Supporting information

**S1 Fig. Flowchart of the SNPs identification methodology using nuclear and mitochondrial genomes sequences**.

(TIFF)

**S2 Fig. Dot-plot analyses comparing three *Euterpe* chloroplast genomes**. (A) *Euterpe edulis* (X – axis) x *Euterpe Oleracea* (Y-axis), (B) *E. edulis* (X-axis) x *Euterpe precatoria* (Y-axis), (C) *E. oleracea* (X – axis) x *E. precatoria* (Y-axis). The positive slope, in purple, represents the pair of sequences aligned and in the same orientation. The negative slope, in blue, represents the pair of sequence aligned, but in opposite orientation. The blue arrow in A highlights the region with inversions and SNPs between *E. edulis* and *E. oleracea*. (D) Local alignment with the chloroplast genomes of *E. edulis, E. oleracea* and *E. precatoria* in the region where inversions and SNPs were detected in A.

(TIFF)

**S3 Fig. Synteny and divergence in the size of the SSC detected in Arecaceae chloroplast genome sequences**. A sample of 20 different chloroplast genomes is shown. Color bars indicate syntenic blocks and the lines indicate the correspondence between them. Blocks on the top row are in the same orientation, while blocks on the bottom row are in inverse orientation. (TIFF)

**Fig S4. SSR and dispersed repeats in the chloroplast genomes of *Euterpe edulis, E. oleracea* and *E. precatoria***. (A) The number of SSR motifs found in the three chloroplast genomes, considering sequence complementarities; (B) Number of dispersed repeats present in different size classes.

(TIFF)

**S5 Fig. Number and type of SNPs found in the alignment of three *Euterpe* chloroplast genomes**

(TIFF)

**S6 Fig. Description of 1**,**077 SNPs found among the nuclear and mitochondrial genome sequences from three *Euterpe* species**. A) Mean depth per loci; B) Mean depth per species sequenced; C) Mutation count in the present in the 1,077 SNPS. (TIFF)

**S7 Fig. Pairwise *F***_***ST***_ **based on 1**,**077 SNPs of nuclear and mitochondrial genomes sequences from three *Euterpe* specie**

(TIFF)

**S1 Table. List of complete chloroplast genomes used in comparative analysis. Synteny and rearrangement, border comparison and phylogenomics**. (XSLX)

**S2 Table. Insertions identified in the alignment of *Euterpe edulis* and *E. oleracea* chloroplast genomes with sequences from GenBank (de Santana Lopes et al. 2021)**. CDS = Coding sequence; IGS = intergenic spacer; LSC = Large single copy; SSC = Small single copy.

(XSLX)

**S3 Table. Simple sequence repeats (SSR) identified in the complete chloroplast genomes of three *Euterpe* species**. CDS = Coding sequence; IGS = intergenic spacer; LSC = Large single copy; SSC = Small single copy; IRs = Inverted Repeats.

(XSLX)

**S4 Table. Simple sequence repeats (SSR) among the chloroplast genomes of *Euterpe edulis, E. oleracea* and *E. precatoria* according to their location**. (XSLX)

**S5 Table. Repeats identified in the chloroplast genomes of *Euterpe edulis, E. oleracea* and *E. precatoria***. F = Forward; P = Palindrome; R = Reverse; C = Complement; CDS= Coding sequence; IGS = intergenic spacer; LSC = Large single copy; SSC = Small single copy; IRs = Inverted Repeats.

(XSLX)

**S6 Table. Indels identified in the pairwise comparison of three *Euterpe* species complete chloroplast genomes**. CDS = Coding sequence; IGS = intergenic spacer; LSC = Large single copy; SSC = Small single copy; IRs = Inverted Repeats. (XSLX)

**S7 Table. Single nucleotide polymorphisms (SNPs) identified in the pairwise comparison of three *Euterpe* species complete chloroplast genomes**. CDS = Coding sequence; IGS = intergenic spacer; LSC = Large single copy; SSC = Small single copy; IRs = Inverted Repeats.

(XSLX)

**S8 Table. Bayesian selection results based on synonymous (Ka)/non-synonymous (Ks) ratio of amino acid substitutions from the *atpE* gene in 54 palm species**. (XSLX)

**S9 Table. Bayesian selection results based on synonymous (Ka)/non-synonymous (Ks) ratio of amino acid substitutions from the *psbJ* gene in 54 palm species**. (XSLX)

## Author Contributions

**Conceptualization:** Maria Imaculada Zucchi

**Data curation:** Ana Flávia Francisconi, Jonathan Andre Morales Marroquín, Cássio van den Berg, Luciano Delmondes de Alencar, Alessandro Alves-Pereira

**Formal Analysis:** Ana Flávia Francisconi, Luiz Augusto Cauz dos Santos, Cássio van den Berg, Alessandro Alves-Pereira, Luciano Delmondes de Alencar

**Funding Acquisition:** Maria Imaculada Zucchi

**Investigation:** Ana Flávia Francisconi, Luiz Augusto Cauz dos Santos

**Methodology:** Luiz Augusto Cauz dos Santos, Jonathan Andre Morales Marroquín, Doriane Picanço-Rodrigues, Luciano Delmondes de Alencar, Cesar Augusto Zanello, Marcones Ferreira Costa

**Resources:** Maria Imaculada Zucchi, Doriane Picanço-Rodrigues

**Software:** Cássio van den Berg

**Supervision:** Maria Imaculada Zucchi

**Writing – Original Draft Preparation:** Ana Flávia Francisconi, Luiz Augusto Cauz dos Santos

**Writing – Review & Editing:** Ana Flávia Francisconi, Luiz Augusto Cauz dos Santos, Jonathan Andre Morales Marroquín, Cássio van den Berg, Alessandro Alves-Pereira, Doriane Picanço-Rodrigues, Luciano Delmondes de Alencar, Cesar Augusto Zanello, Marcones Ferreira Costa, Maria Teresa Gomes Lopes, Elizabeth Ann Veasey, Maria Imaculada Zucchi

## References

1. Uthaipaisanwong P, Chanprasert J, Shearman JR, Sangsrakru D, Yoocha T, Jomchai N, et al. Characterization of the chloroplast genome sequence of oil palm (Elaeis guineensis Jacq.). Gene. 2012;500: 172–180. doi:10.1016/j.gene.2012.03.061

2. Baker WJ, Couvreur TLP. Global biogeography and diversification of palms sheds light on the evolution of tropical lineages. I. Historical biogeography. J Biogeogr. 2013;40: 274–285. doi:10.1111/j.1365-2699.2012.02795.x

3. Brokamp G, Valderrama N, Mittelbach M, Grandez CA, Barfod AS, Weigend M. Trade in palm products in North-Western South America. Bot Rev. 2011;77: 571–606. doi:10.1007/s

4. Eiserhardt WL, Pintaud J, Asmussen-lange C, Hahn J, Bernal R, Balslev H, et al. Phytogeny and divergence times of Bactridinae (Arecaceae, Palmae) based on plastid and nuclear DNA sequences. 2011;60: 485–498.

5. Baker WJ, Couvreur TLP. Global biogeography and diversification of palms sheds light on the evolution of tropical lineages. II. Diversification history and origin of regional assemblages. J Biogeogr. 2013;40: 286–298. doi:10.1111/j.1365-2699.2012.02794.x

6. Henderson A, Galeano G. Euterpe, Prestoea, and Neonicholsonia (Palmae). Flora Neotrop. 1996;72: 1–89.

7. Mart E, Bussmann RW, Zambrana NYP. Acta Societatis Botanicorum Poloniae Facing global markets – usage changes in Western Amazonian plants: the example of Euterpe precatoria Mart. and E. oleracea Mart. 2012;81: 257–261. doi:10.5586/asbp.2012.032

8. Galetti M, Guevara R, Côrtes MC, Fadini R, Matter S Von, Leite AB, et al. Functional extinction of birds drives rapid evolutionary changes in seed size. Science (80-). 2013;340: 1086–1091. doi:DOI: 10.1126/science.1233774

9. Alves-Pereira A, Novello M, Dequigiovanni G, Pinheiro JB, Brancalion PHS, Veasey EA, et al. Genomic diversity of three Brazilian native food crops based on double-digest restriction site-associated DNA sequencing. Trop Plant Biol. 2019;12: 268–281. doi:10.1007/s12042-019-09229-z

10. Novello M, Viana JPG, Alves-Pereira A, de Aguiar Silvestre E, Nunes HF, Pinheiro JB, et al. Genetic conservation of a threatened Neotropical palm through community-management of fruits in agroforests and second-growth forests. For Ecol Manage. 2017;407: 200–209. doi:10.1016/j.foreco.2017.06.059

11. Coelho GM, Santos AS, de Menezes IPP, Tarazi R, Souza FMO, Silva M das GCPC, et al. Genetic structure among morphotypes of the endangered Brazilian palm Euterpe edulis Mart (Arecaceae). Ecol Evol. 2020;10: 6039–6048. doi:10.1002/ece3.6348

12. Martinelli G, Moraes MA. Livro vermelho da flora do Brasil. 1st ed. Rio de Janeiro: CNCFlora, Centro Nacional de Conservação da Flora Rio de Janeiro; 2013.

13. Ball AA, Brancalion PHS. Governance challenges for commercial exploitation of a non-timber forest product by marginalized rural communities. Environ Conserv. 2016;43: 208–220. doi:10.1017/S0376892916000072

14. IBGE. Censo agropecuário 2017: resultados definitivos. Censo agropecuário. 2019;8: 1–105. Available from: https://biblioteca.ibge.gov.br/visualizacao/periodicos/3096/agro_2017_resultados_definitivos.pdf

15. Ramos SLF, Dequigiovanni G, Lopes MTG, Aguiar AV de, Lopes R, Veasey EA, et al. Genetic structure in populations of Euterpe precatoria Mart. in the Brazilian Amazon. Front Ecol Evol. 2021;8: 1–11. doi:10.3389/fevo.2020.603448

16. Shanley P, Medina G. Frutíferas e plantas úteis na vida amazônica. 1st ed. Belém-PA: Cifor, Imazon; 2005.

17. Vedel-sørensen M, Tovaranonte J, Bøcher PK, Balslev H, Barfod AS. Spatial distribution and environmental preferences of 10 economically important forest palms in western South America. For Ecol Manage. 2013;307: 284–292. doi:10.1016/j.foreco.2013.07.005

18. Freitas MAB, Vieira ICG, Albernaz ALKM, Magalhães JLL, Lees AC. Floristic impoverishment of Amazonian floodplain forests managed for açaí fruit production. For Ecol Manage. 2015;351: 20–27. doi:10.1016/j.foreco.2015.05.008

19. Campbell AJ, Carvalheiro LG, Maués MM, Jaffé R, Giannini TC, Freitas MAB, et al. Anthropogenic disturbance of tropical forests threatens pollination services to açaí palm in the Amazon river delta. J Appl Ecol. 2018; 1725–1736. doi:10.1111/1365-2664.13086

20. Kahn F. Palms as key swamp forest resources in Amazonia. For Ecol Manage. 1991;38: 133–142. doi:https://doi.org/10.1016/0378-1127(91)90139-M

21. Ramos SLF, Dequigiovanni G, Magno A, Pimentel P, Teresa M, Lopes G, et al. Paternity analysis, pollen flow, and spatial genetic structure of a natural population of Euterpe precatoria in the Brazilian Amazon. Ecol Evol. 2018;8: 11143–11157. doi:10.1002/ece3.4582

22. Du FK, Lang T, Lu S, Wang Y, Li J, Yin K. An improved method for chloroplast genome sequencing in non-model forest tree species. Tree Genet Genomes. 2015;11. doi:10.1007/s11295-015-0942-2

23. Dodsworth S. Genome skimming for next-generation biodiversity analysis. Trends Plant Sci. 2015;20: 525–527. doi:10.1016/j.tplants.2015.06.012

24. Sobreiro MB, Vieira LD, Nunes R, Novaes E, Coissac E, Silva-Junior OB, et al. Chloroplast genome assembly of Handroanthus impetiginosus: Comparative analysis and molecular evolution in Bignoniaceae. Planta. 2020;252: 1–16. doi:10.1007/s00425-020-03498-9

25. Sugiura M. The chloroplast genome. Plant Mol Biol. 1992;19: 149–168. doi:10.1007/978-94-011-2656-4_10

26. Barrett CF, Bacon CD, Antonelli A, Cano Á, Hofmann T. An introduction to plant phylogenomics with a focus on palms. Bot J Linn Soc. 2016;182: 234–255. doi:10.1111/boj.12399

27. Dabral A, Shamoon A, Meena RK, Kant R, Pandey S, Ginwal HS, et al. Genome skimming-based simple sequence repeat (SSR) marker discovery and characterization in Grevillea robusta. Physiol Mol Biol Plants. 2021;27: 1623–1638. doi:10.1007/s12298-021-01035-w

28. de Santana Lopes A, Gomes Pacheco T, Nimz T, do Nascimento Vieira L, Guerra MP, Nodari RO, et al. The complete plastome of macaw palm [Acrocomia aculeata (Jacq.) Lodd. ex Mart.] and extensive molecular analyses of the evolution of plastid genes in Arecaceae. Planta. 2018;247: 1011–1030. doi:10.1007/s00425-018-2841-x

29. Aljohi HA, Liu W, Lin Q, Zhao Y, Zeng J, Alamer A, et al. Complete sequence and analysis of coconut palm (Cocos nucifera) mitochondrial genome. PLoS One. 2016;11: 1–18. doi:10.1371/journal.pone.0163990

30. Yang Y, Zhou T, Duan D, Yang J, Feng L, Zhao G. Comparative analysis of the complete chloroplast genomes of five quercus species. Front Plant Sci. 2016;7: 1–13. doi:10.3389/fpls.2016.00959

31. Cauz-santos LA, Munhoz CF, Rodde N, Cauet S, Santos AA, Penha HA, et al. The chloroplast genome of Passiflora edulis (Passifloraceae) assembled from long sequence reads: structural organization and phylogenomic studies in malpighiales. 2017;8: 1–17. doi:10.3389/fpls.2017.00334

32. Machado L de O, Vieira L do N, Stefenon VM, Oliveira Pedrosa F de, Souza EM de, Guerra MP, et al. Phylogenomic relationship of feijoa (Acca sellowiana (O.Berg) Burret) with other Myrtaceae based on complete chloroplast genome sequences. Genetica. 2017;145: 163–174. doi:10.1007/s10709-017-9954-1

33. Pichardo-Marcano FJ, Nieto-Blázquez ME, MacDonald AN, Galeano G, Roncal J. Phylogeny, historical biogeography and diversification rates in an economically important group of Neotropical palms: Tribe Euterpeae. Mol Phylogenet Evol. 2019;133: 67–81. doi:10.1016/j.ympev.2018.12.030

34. de Santana Lopes A, Gomes Pacheco T, Nascimento da Silva O, do Nascimento Vieira L, Guerra MP, Pacca Luna Mattar E, et al. Plastid genome evolution in Amazonian açaí palm (Euterpe oleracea Mart.) and Atlantic forest açaí palm (Euterpe edulis Mart.). Plant Mol Biol. 2021;105: 559–574. doi:10.1007/s11103-020-01109-5

35. Liu L, Zhang Y, Li P. Development of genomic resources for the genus Celtis (Cannabaceae) based on genome skimming data. Plant Divers. 2021;43: 43–53. doi:10.1016/j.pld.2020.09.005

36. Takamatsu T, Baslam M, Inomata T, Oikawa K. Optimized method of extracting rice chloroplast DNA for high-quality plastome resequencing and de novo assembly. 2018;9: 1–13. doi:10.3389/fpls.2018.00266

37. Dierckxsens N, Mardulyn P, Smits G. NOVOPlasty: De novo assembly of organelle genomes from whole genome data. Nucleic Acids Res. 2017;45. doi:10.1093/nar/gkw955

38. Jin JJ, Yu W Bin, Yang JB, Song Y, DePamphilis CW, Yi TS, et al. GetOrganelle: A fast and versatile toolkit for accurate de novo assembly of organelle genomes. Genome. 2020;21: 1–31. doi:10.1101/256479

39. Langdon WB. Performance of genetic programming optimised Bowtie2 on genome comparison and analytic testing (GCAT) benchmarks. BioData Min. 2015;8: 1–7. doi:10.1186/s13040-014-0034-0

40. Bankevich A, Nurk S, Antipov D, Gurevich AA, Dvorkin M, Kulikov AS, et al. SPAdes: A new genome assembly algorithm and its applications to single-cell sequencing. J Comput Biol. 2012;19: 455–477. doi:10.1089/cmb.2012.0021

41. Tillich M, Lehwark P, Pellizzer T, Ulbricht-Jones ES, Fischer A, Bock R, et al. GeSeq - Versatile and accurate annotation of organelle genomes. Nucleic Acids Res. 2017;45: W6–W11. doi:10.1093/nar/gkx391

42. Abeel T, Van Parys T, Saeys Y, Galagan J, Van De Peer Y. GenomeView: A next-generation genome browser. Nucleic Acids Res. 2012;40: 1–10. doi:10.1093/nar/gkr995

43. Greiner S, Lehwark P, Bock R. OrganellarGenomeDRAW (OGDRAW) version 1.3.1: Expanded toolkit for the graphical visualization of organellar genomes. Nucleic Acids Res. 2019;47: W59–W64. doi:10.1093/nar/gkz238

44. Marçais G, Delcher AL, Phillippy AM, Coston R, Salzberg SL, Zimin A. MUMmer4: A fast and versatile genome alignment system. PLoS Comput Biol. 2018;14: 1–14. doi:10.1371/journal.pcbi.1005944

45. Katoh K, Standley DM. MAFFT: Iterative refinement and additional methods. Methods Mol Biol. 2014;1079: 131–146. doi:10.1007/978-1-62703-646-7_8

46. Da Silva RS, Clementi CR, Balsanelli E, de Baura VA, de Souza EM, de Freitas Fraga HP, et al. The plastome sequence of Bactris gasipaes and evolutionary analysis in tribe Cocoseae (Arecaceae). PLoS One. 2021;16: 1–15. doi:10.1371/journal.pone.0256373

47. Darling ACE, Mau B, Blattner FR, Perna NT. Mauve: Multiple alignment of conserved genomic sequence with rearrangements. Genome Res. 2004;14: 1394–1403. doi:10.1101/gr.2289704

48. Beier S, Thiel T, Münch T, Scholz U, Mascher M. MISA-web: A web server for microsatellite prediction. Bioinformatics. 2017;33: 2583–2585. doi:10.1093/bioinformatics/btx198

49. Kurtz S, Schleiermacher C. REPuter: fast computation of maximal repeats in complete genomes. Bioinforma Appl Note. 1999;15: 426–427. doi:10.1093/bioinformatics/15.5.426

50. Librado P, Rozas J. DnaSP v5: A software for comprehensive analysis of DNA polymorphism data. Bioinformatics. 2009;25: 1451–1452. doi:10.1093/bioinformatics/btp187

51. Edgar RC. MUSCLE: Multiple sequence alignment with high accuracy and high throughput. Nucleic Acids Res. 2004;32: 1792–1797. doi:10.1093/nar/gkh340

52. Stern A, Doron-Faigenboim A, Erez E, Martz E, Bacharach E, Pupko T. Selecton 2007: Advanced models for detecting positive and purifying selection using a Bayesian inference approach. Nucleic Acids Res. 2007;35: 506–511. doi:10.1093/nar/gkm382

53. Nylander JAA. MrModeltest v2. Program distributed by the author. Evolutionary Biology Center, Uppsala University; 2004.

54. Xie W, Lewis PO, Fan Y, Kuo L, Chen MH. Improving marginal likelihood estimation for bayesian phylogenetic model selection. Syst Biol. 2011;60: 150–160. doi:10.1093/sysbio/syq085

55. Ronquist F, Teslenko M, Van Der Mark P, Ayres DL, Darling A, Höhna S, et al. Mrbayes 3.2: Efficient bayesian phylogenetic inference and model choice across a large model space. Syst Biol. 2012;61: 539–542. doi:10.1093/sysbio/sys029

56. Miller MA, Pfeiffer W, Schwartz T. Creating the CIPRES Science Gateway for inference of large phylogenetic trees. 2010 Gatew Comput Environ Work GCE 2010. 2010. doi:10.1109/GCE.2010.5676129

57. Kass RE, Raftery AE. Bayes Factors. J Am Stat Assoc. 1995;90: 773–795.

58. Rambaut A. FigTree v1. 3.1. 2012. Available from: http://tree.bio.ed.ac.uk/software/figtree

59. Inkscape. Inkscape Project. 2021. Available from: https://inkscape.org/pt-br/

60. Li H, Handsaker B, Wysoker A, Fennell T, Ruan J, Homer N, et al. The Sequence Alignment/Map format and SAMtools. Bioinformatics. 2009;25: 2078–2079. doi:10.1093/bioinformatics/btp352

61. Broad Institute. Picard Tools. Broad Institute GitHub Repository. 2018. Available from: http://broadinstitute.github.io/picard/

62. Catchen J, Hohenlohe PA, Bassham S, Amores A. Stacks: an analysis tool set for population genomics. 2013; 3124–3140. doi:10.1111/mec.12354

63. Danecek P, Auton A, Abecasis G, Albers CA, Banks E, DePristo MA, et al. The variant call format and VCFtools. Bioinformatics. 2011;27: 2156–2158. doi:10.1093/bioinformatics/btr330

64. Jombart T, Ahmed I. adegenet 1.3-1: New tools for the analysis of genome-wide SNP data. Bioinformatics. 2011;27: 3070–3071. doi:10.1093/bioinformatics/btr521

65. Rousset F, Lopez L, Belkhir K. R package: genepop. 2020. p. 16. Available from: http://kimura.univ-montp2.fr/~rousset/Genepop.htm

66. R Core Team. R: A language and environment for statistical computing. Vienna, Austria: R Foundation for Statistical Computing; 2021. Available from: https://www.r-project.org/

67. de Santana Lopes A, Gomes Pacheco T, Nascimento da Silva O, Magalhães Cruz L, Balsanelli E, Maltempi de Souza E, et al. The plastomes of Astrocaryum aculeatum G. Mey. and A. murumuru Mart. show a flip-flop recombination between two short inverted repeats. Planta. 2019;250: 1229–1246. doi:10.1007/s00425-019-03217-z

68. Drescher A, Stephanie R, Calsa T, Carrer H, Bock R. The two largest chloroplast genome-encoded open reading frames of higher plants are essential genes. Plant J. 2000;22: 97–104. doi:10.1046/j.1365-313X.2000.00722.x

69. Kikuchi S, Asakura Y, Imai M, Nakahira Y, Kotani Y, Hashiguchi Y, et al. A Ycf2-FtsHi heteromeric AAA-ATPase complex is required for chloroplast protein import. Plant Cell. 2018;30: 2677–2703. doi:10.1105/tpc.18.00357

70. Barrett CF, Davis JI, Leebens-Mack J, Conran JG, Stevenson DW. Plastid genomes and deep relationships among the commelinid monocot angiosperms. Cladistics. 2013;29: 65–87. doi:10.1111/j.1096-0031.2012.00418.x

71. Vieira MLC, Santini L, Diniz AL, Munhoz C de F. Microsatellite markers: What they mean and why they are so useful. Genet Mol Biol. 2016;39: 312–328. doi:10.1590/1678-4685-GMB-2016-0027

72. Eguiluz M, Rodrigues NF, Guzman F, Yuyama P, Margis R. The chloroplast genome sequence from Eugenia uniflora, a Myrtaceae from Neotropics. Plant Syst Evol. 2017;303: 1199–1212. doi:10.1007/s00606-017-1431-x

73. Ping J, Feng P, Li J, Zhang R, Su Y, Wang T. Molecular evolution and SSRs analysis based on the chloroplast genome of Callitropsis funebris. Ecol Evol. 2021;11: 4786–4802. doi:10.1002/ece3.7381

74. Rogalski M, Vieira LDN, Fraga HP, Guerra MP. Plastid genomics in horticultural species: Importance and applications for plant population genetics, evolution, and biotechnology. Front Plant Sci. 2015;6: 1–17. doi:10.3389/fpls.2015.00586

75. Zhu A, Guo W, Gupta S, Fan W, Mower JP. Evolutionary dynamics of the plastid inverted repeat: The effects of expansion, contraction, and loss on substitution rates. New Phytol. 2016;209: 1747–1756. doi:10.1111/nph.13743

76. Zeng S, Zhou T, Han K, Yang Y, Zhao J, Liu ZL. The complete chloroplast genome sequences of six rehmannia species. Genes (Basel). 2017;8. doi:10.3390/genes8030103

77. Milligan BG, Hampton JN, Palmer JD. Dispersed repeats and structural reorganization in subclover chloroplast DNA. Mol Biol Evol. 1989;6: 355–368. doi:10.1093/oxfordjournals.molbev.a040558

78. Song Y, Dong W, Liu B, Xu C, Yao X, Gao J, et al. Comparative analysis of complete chloroplast genome sequences of two tropical trees Machilus yunnanensis and Machilus balansae in the family Lauraceae. Front Plant Sci. 2015;6: 1–8. doi:10.3389/fpls.2015.00662

79. Barrett CF, Baker WJ, Comer JR, Conran JG, Lahmeyer SC, Leebens-Mack JH, et al. Plastid genomes reveal support for deep phylogenetic relationships and extensive rate variation among palms and other commelinid monocots. New Phytol. 2016;209: 855–870. doi:10.1111/nph.13617

80. Brancalion PHS, Oliveira GCX, Zucchi MI, Novello M, van Melis J, Zocchi SS, et al. Phenotypic plasticity and local adaptation favor range expansion of a Neotropical palm. Ecol Evol. 2018;8: 7462–7475. doi:10.1002/ece3.4248

81. Du X, Zeng T, Feng Q, Hu L, Luo X, Weng Q, et al. The complete chloroplast genome sequence of yellow mustard (Sinapis alba L.) and its phylogenetic relationship to other Brassicaceae species. Gene. 2020;731. doi:10.1016/j.gene.2020.144340

82. Liu Q, Li X, Li M, Xu W, Schwarzacher T, Heslop-Harrison JS. Comparative chloroplast genome analyses of Avena: Insights into evolutionary dynamics and phylogeny. BMC Plant Biol. 2020;20: 1–20. doi:10.1186/s12870-020-02621-y

83. Huang Y, Yang Z, Huang S, An W, Li J, Zheng X. Comprehensive analysis of Rhodomyrtus tomentosa chloroplast genome. Plants. 2019;8. doi:10.3390/plants8040089

84. Williams-Carrier R, Brewster C, Belcher SE, Rojas M, Chotewutmontri P, Ljungdahl S, et al. The Arabidopsis pentatricopeptide repeat protein LPE1 and its maize ortholog are required for translation of the chloroplast psbJ RNA. Plant J. 2019;99: 56–66. doi:10.1111/tpj.14308

85. Díaz BG, Zucchi MI, Alves-Pereira A, de Almeida CP, Moraes ACL, Vianna SA, et al. Genome-wide SNP analysis to assess the genetic population structure and diversity of Acrocomia species. PLoS One. 2021;16: 1–24. doi:10.1371/journal.pone.0241025

86. Viana JPG, Siqueira MVBM, Araujo FL, Grando C, Sujii PS, De Aguiar Silvestre E, et al. Genomic diversity is similar between Atlantic Forest restorations and natural remnants for the native tree Casearia sylvestris Sw. PLoS One. 2019;13: 1–14. doi:10.1371/journal.pone.0192165

87. Cordeiro EMG, Macrini CM, Sujii PS, Schwarcz KD, Pinheiro JB, Rodrigues RR, et al. Diversity, genetic structure, and population genomics of the tropical tree Centrolobium tomentosum in remnant and restored Atlantic forests. Conserv Genet. 2019;20: 1073–1085. doi:10.1007/s10592-019-01195-z

88. Allendorf FW, Hohenlohe PA, Luikart G. Genomics and the future of conservation genetics. Nat Rev Genet. 2010;11: 697–709. doi:10.1038/nrg2844

89. Willing EM, Dreyer C, van Oosterhout C. Estimates of genetic differentiation measured by fst do not necessarily require large sample sizes when using many snp markers. PLoS One. 2012;7: 1–7. doi:10.1371/journal.pone.0042649

90. Hamblin MT, Warburton ML, Buckler ES. Empirical comparison of simple sequence repeats and single nucleotide polymorphisms in assesment of maize diversity and relatedness. PLoS One. 2007;2. doi:10.1371/journal.pone.0001367

